# Hierarchical taxonomic constraints shape bacterial nitrogen cycling across functional and molecular scales

**DOI:** 10.1101/2025.11.26.690687

**Authors:** Mingli Jiang, Yanni Huang, Zhiming Wu, Qian Zhu, Qian Li, Kaihua Pan, Mingliang Zhang, Liang Shi, Jiguo Qiu, Pengfa Li, Yiyong Zhu, Xin Yan, Qing Hong

**Affiliations:** Department of Microbiology, College of Life Sciences, Nanjing Agricultural University, Key Laboratory of Agricultural and Environmental Microbiology, Ministry of Agriculture and Rural Affairs, Nanjing 210095, China; College of Resources and Environment Sciences, Nanjing Agricultural University, Nanjing 210095, China; College of Resources and Environment, Fujian Agriculture and Forestry University, Fuzhou 350002, China; College of Life Sciences, Nanjing Agricultural University, Nanjing 210095, China; Nanchang Key Laboratory of Microbial Resources Exploitation & Utilization from Poyang Lake Wetland, College of Life Sciences, Jiangxi Normal University, Nanchang 330022, China

**Author notes:** Author for correspondence: Qing Hong E-mail address., Physical mailing address: Qing Hong, College of Life Sciences, Nanjing Agricultural University, 1 Weigang, Xuanwu District, Nanjing, Jiangsu Province, 210095, China.

**Keywords:** Nitrogen cycling, Taxonomic constraints, Functional archetypes, Comparative genomics, Sequence conservation

## Abstract

Microbes are essential for global nitrogen cycling, yet the extent to which taxonomic identity constrains functional potential remains poorly quantified. Using 73,472 representative bacterial genomes, we establish a multi-scale quantitative framework revealing systematic, hierarchical relationships between taxonomic identity and nitrogen cycling functional potential. Hierarchical variance decomposition reveals that taxonomy explains 20–46% of functional variation across six nitrogen pathways, with the strongest constraints for dissimilatory nitrite reduction to ammonium (46.1%) and nitrogen fixation (44.2%), both negatively correlated with functional prevalence. K-means clustering identifies four class-level functional archetypes (Functionally Inactive, Diversified, Integrated, and Specialized) among 77 bacterial classes. 1,281 genera resolve into five ecological strategies differentiated by their nitrogen retention versus loss capabilities, exhibiting strong phylogenetic signals. Cross-scale validation demonstrates 78.7% genus-level conformity to class-level archetypes, confirming hierarchical functional organization. Molecular evolutionary analysis of 13 genes reveals sequence conservation as a dimension partially independent of pathway-level functional constraints. Paradoxically, the functionally most constrained pathway exhibits only moderate sequence conservation (*nrfA*: 9th/13 genes), while a moderately constrained pathway shows exceptional conservation (*napA*). Four constraint-conservation patterns demonstrate that gene-specific structural and ecological factors generate evolutionary rate variation independently of taxonomic associations. Our results establish a hierarchical framework in which taxonomic constraints set baseline functional potential, ecological trade-offs shape strategy diversification, and molecular evolution modulates gene-level conservation patterns across biological scales. This framework establishes quantitative baselines that enable probabilistic inference of nitrogen cycling capabilities from taxonomic composition, with potential applications in amplicon-based community analysis, targeted cultivation, and biogeochemical modeling.

**Importance:** A fundamental challenge in microbial ecology is inferring functional potential from taxonomic data—a relationship widely assumed but never rigorously quantified. Resolving this is critical because while amplicon sequencing provides cost-effective taxonomic profiling, functional characterization requires expensive metagenomics, limiting large-scale biogeochemical studies. We provide the first systematic quantification demonstrating that taxonomic identity explains 20-46% of nitrogen cycling functional variation, operating hierarchically from class to genus level. Crucially, we reveal that taxonomic constraints operate at both functional distribution and molecular evolution levels as partially independent dimensions, indicating distinct evolutionary mechanisms. This work establishes quantitative foundations for taxonomy-based functional prediction, enabling researchers to extract functional insights from readily available taxonomic surveys. As reference databases expand, this framework will enhance predictive capabilities for nitrogen cycling and broader biogeochemical processes.

## Introduction

Nitrogen cycling controls ecosystem productivity and represents one of the most important Earth system functions, with bacterial microbial communities driving nitrogen transformations through six principal pathways: nitrogen fixation (NF), nitrification (NIT), denitrification (DNF), dissimilatory nitrate reduction (DNN), assimilatory nitrate reduction (ANRA), and dissimilatory nitrite reduction to ammonium (DNRA) (Figure S1) (1, 2). These processes collectively govern nitrogen retention, loss, and bioavailability across ecosystems, fundamentally influencing primary productivity and greenhouse gas emissions (3). Understanding the distribution of nitrogen cycling capabilities across bacterial diversity therefore represents a foundational challenge for predicting ecosystem responses and managing nitrogen pollution in the context of global environmental change (4, 5).

Taxonomic identity constitutes the most fundamental information in microbial studies, yet quantitative relationships between taxonomic lineages and nitrogen cycling functional potential remain poorly characterized. Bacterial genera exhibit striking variation in nitrogen cycling repertoires, ranging from specialists like *Pectobacterium* (primarily encoding nitrogen fixing genes) to generalists like *Anaeromyxobacter* (harboring most nitrogen cycling genes) (6). While phylogenetic conservatism of microbial traits has been demonstrated (7, 8), the strength of taxonomic associations with nitrogen cycling functional potential, and how these associations vary among transformation pathways, has not been systematically quantified. Nitrogen cycling capabilities are shaped by both vertical inheritance within taxonomic lineages and horizontal gene transfer (HGT) across phylogenetically distant taxa (9). The relative contributions of these evolutionary mechanisms to functional distribution patterns remain unquantified, and whether different pathways exhibit distinct constraint strengths due to varying evolutionary pressures and pathway complexity (10), has not been systematically assessed across bacterial diversity.

Existing studies have been limited in scale and scope. Previous investigations of nitrogen cycling functions, whether using comparative genomics or environmental metagenomics approaches, have typically examined hundreds to a few thousand genomes and focused on specific pathways or limited environmental contexts (6, 11). While these studies have successfully documented the presence and distribution of nitrogen cycling genes across microbial diversity, they rarely quantify the strength of taxonomic associations with functional potential or systematically compare constraint patterns across multiple transformation pathways. Moreover, recent work has documented phylogenetic signals (7) and taxonomic patterns (6), but has not quantified constraint strength or established correlations with pathway-level properties. The relationship between functional prevalence and taxonomic constraint strength also remains largely unexplored, despite evidence that widely distributed functions may exhibit weaker phylogenetic conservation than complex, multi-gene traits (12, 13). Addressing these gaps requires a large-scale, systematic analysis that quantifies constraint patterns across multiple pathways and links these patterns to pathway-specific evolutionary properties.

The Genome Taxonomy Database (GTDB) now provides unprecedented opportunities for large-scale analysis, encompassing hundreds of thousands of bacterial genomes with consistent taxonomic assignments (14). Hierarchical variance decomposition methods can quantify taxonomic association strength using taxonomic structure as a proxy for evolutionary relationships (15). This approach enables the first comprehensive quantitative framework for understanding how taxonomic identity relates to nitrogen cycling functional potential across bacterial diversity.

Here, we address how taxonomic identity constrains nitrogen cycling functional potential across bacterial diversity. Using 73,472 representative bacterial genomes, we develop a multi-scale quantitative framework integrating hierarchical variance decomposition, unsupervised functional classification, and molecular evolutionary analysis to address three specific questions. First, can taxonomic constraints be quantified, and do constraint strengths vary systematically across pathways in ways that reflect pathway-level properties? Second, does functional organization exhibit hierarchical structure across taxonomic scales, with class-level patterns constraining genus-level ecological strategies? Third, do pathway-level functional constraints translate uniformly into molecular-level sequence conservation, or do gene-specific factors decouple these evolutionary dimensions? Collectively, these analyses will establish quantitative baselines for assessing taxonomy-function associations in bacterial nitrogen cycling, providing empirical foundations for taxonomy-based functional inference.

## Materials and Methods

### Data sources and annotation pipeline

High-quality bacterial genomes were obtained from the Genome Taxonomy Database (GTDB) release 220 (14). Genomes were selected based on GTDB representative status or type species designation with quality control criteria of completeness >90% and contamination <10%. The final dataset comprised 73,472 bacterial genomes.

Functional annotation employed a two-step integrated approach. Primary annotation was performed using KEGG database (16) and RAST server (17) to identify all potential nitrogen metabolism-related genes. Genes identified as potentially involved in nitrogen cycling were subsequently annotated using the specialized NCycDB database (18) with DIAMOND (19) using parameters ‘-k 1-e 0.0001’ (20). Final functional assignment required concordant annotation from at least two of the three databases, providing a majority-vote consensus to resolve potential conflicts. This hierarchical strategy combines comprehensive pathway screening with specialized nitrogen cycling gene annotation to reduce both false negatives and false positives. Annotation quality was validated using 30 single-copy housekeeping genes as internal controls (6), with consistent detection (>95% recovery rate) confirming pipeline reliability.

### Functional gene criteria and environmental classification

To determine nitrogen cycling functional potential across bacterial genomes, gene selection focused on genes encoding enzymes catalyzing key steps in nitrogen transformation pathways (Figure S1). Functional potential was assessed using a dual-detection strategy (6, 11) where a genome was assigned functional capability only when all required core genes were detected alongside at least one auxiliary gene for that pathway. Core and auxiliary genes for six pathways were identified from previous studies (1, 6, 11) (Supplementary Table S1 and Supplementary EM1). Core-auxiliary gene correlation analysis validated significant associations between core and auxiliary genes within pathways (Figure S2).

Environmental metadata were obtained from NCBI BioSample, documenting the original sampling environment from which each genome was obtained. Genomes were classified into six broad environmental categories using standardized keywords: Host-associated, Aquatic, Terrestrial, Extreme, Engineered, and Other (21). These categories reflect the genome source environment—the physical location or substrate from which the biological sample was originally collected—encompassing both cultivated isolates (from environmental enrichments or direct isolation) and metagenome-assembled genomes (MAGs) (from environmental DNA sequencing).

Classification accuracy was validated through manual verification of 300 randomly selected samples (50 per category, κ = 0.92, 95% CI: 0.89-0.95) (22). For higher-resolution analysis, environmental sources were further subdivided into 22 detailed subtypes based on refined metadata extraction (Supplementary EM2). Genome source categories represent the sampling origin of sequenced genomes rather than strict ecological niches. While public genome databases are subject to well-documented biases—including cultivation feasibility, geographic sampling intensity, and research priorities (23, 24)—the observed environmental associations nonetheless reflect biogeographical patterns in bacterial diversity that are consistent with established ecological and biogeochemical principles, as validated in the Results section.

### Quantifying taxonomic associations with functional potential

Taxonomic associations were quantified using hierarchical variance decomposition, with taxonomic structure as a proxy for evolutionary relationships given computational constraints at this scale. This approach measures vertical inheritance patterns rather than direct phylogenetic constraints (Supplementary EM3). Generalized linear mixed-effects models (GLMM) with binomial family and logit link were applied using lme4 package (25) to quantify the strength of taxonomic associations with functional potential. Analysis was restricted to genera with ≥10 genomes and classes with ≥3 genera and ≥50 genomes (26).

Taxonomic structure was modeled as nested random effects (Class/Genus) in the model: response ∼ (1|Class/Genus), where functional presence/absence was predicted solely by taxonomic identity. Taxonomic association strength was calculated as σ²class / (σ²class + σ²genus + π²/3), where σ²class and σ²genus represent class-level and genus-level variance components respectively, and π²/3 represents the variance of the logistic distribution (15, 27). This approach quantifies the proportion of total functional variation explained by taxonomic identity, which captures both vertical inheritance patterns and historical evolutionary signals preserved in the taxonomic framework. Methodological rationale and model diagnostics are detailed in Supplementary EM3.

For functions with extreme rarity (NIT: 0.3% prevalence), Fisher’s exact tests were employed to assess taxonomic clustering patterns across bacterial classes. Bootstrap confidence intervals (1000 replicates) were computed where model convergence allowed (Supplementary EM3). Multiple testing correction was applied using Benjamini-Hochberg method across all six nitrogen cycling functions (28).

### Class-level functional strategy classification and environmental modulation

Class-level functional strategies were characterized using participation rate (proportion of genomes with ≥1 nitrogen cycling function) and functional concentration (Simpson concentration index (29). K-means clustering using cluster package (30) identified optimal cluster number (k = 4) through silhouette analysis (31). Analysis included bacterial classes with ≥100 genomes. Environmental modulation and function-environment associations were assessed as detailed in Supplementary EM4.

### Genus-level ecological strategies and evolutionary signal analysis

Genus-level ecological strategy classification employed a five-type framework based on nitrogen biogeochemical function profiles. Genera were classified based on nitrogen retention versus nitrogen loss processes: N-Retention (nitrogen retention functions: ANRA, NF, or DNRA only); N-Loss (nitrogen loss functions: DNF or NIT only); Multifunctional (both retention and loss functions); Metabolically Versatile (DNN function only); Non-N-Cycling (no detectable nitrogen cycling functions). This classification captures biogeochemical trade-offs in microbial nitrogen metabolism (1). Phylogenetic signal analysis, functional heterogeneity quantification, and statistical testing procedures are detailed in Supplementary EM5.

### Molecular evolutionary analysis and multi-scale conservation patterns

Nucleotide diversity analysis was performed on 13 nitrogen cycling genes to assess sequence conservation across multiple biological scales. For each gene, nucleotide sequences were aligned using MAFFT (32), and nucleotide diversity (π) was calculated as the average pairwise nucleotide differences per site (33). Multi-scale nucleotide diversity analysis quantified evolutionary constraints at genus, class, and environmental levels by calculating π separately for genes grouped by each taxonomic or environmental category, where lower π values indicate greater sequence conservation. For π calculations, genera with ≥3 sequences were retained to permit pairwise diversity estimation. Environmental sensitivity was quantified as the coefficient of variation of π values across environmental contexts, with bootstrap confidence intervals (n = 1000) calculated using the boot package (34).

To assess amino acid-level selection pressures, we conducted dN/dS ratio (ω) analyses using codon-based evolutionary models. For dN/dS analysis, only genera with ≥20 sequences were retained to ensure reliable phylogenetic signal strength and codon-based model fitting. Global dN/dS ratios were estimated using the Single-Likelihood Ancestor Counting (SLAC) method implemented in HyPhy (35). We report median ω values with interquartile ranges (IQR) to summarize selection pressures across genera. Differences in nucleotide diversity across taxonomic scales were assessed using paired Wilcoxon signed-rank tests (36). Values of ω < 0.3 indicate strong purifying selection, 0.3 ≤ ω < 1.0 indicate moderate purifying selection, and ω ≥ 1.0 suggest neutral evolution or positive selection. Detailed quality control procedures for sequence alignment, codon integrity, and statistical methods are provided in Supplementary EM6.

### Statistical analysis and data visualization

Functional co-occurrence patterns were assessed using Jaccard similarity indices [51] (J = |A∩B| / |A∪B|), where A and B represent gene sets for pathway pairs. Statistical enrichment of function-environment and taxonomic associations was evaluated using two-tailed Fisher’s exact tests, with odds ratios calculated from 2×2 contingency tables. Enrichment (OR > 1.5) and depletion (OR < 0.67) thresholds were defined a priori based on biological relevance. Cross-scale conformity was quantified as the proportion of genera exhibiting ecological strategies consistent with their class-level functional archetypes.

Unless otherwise specified, statistical significance was assessed at α = 0.05 with Benjamini-Hochberg false discovery rate correction applied for multiple comparisons (28). All statistical analyses were conducted in R version 4.3.0 (37). Data visualization tools and parameters are detailed in Supplementary EM7.

## Results

### Functional sparsity and environment-specific enrichment characterize bacterial nitrogen cycling potential

Nitrogen cycling capabilities are unevenly distributed across bacterial diversity. Analyzing 73,472 representative bacterial genomes from the Genome Taxonomy Database (GTDB), we found pronounced functional sparsity: 54.3% of genomes lacked any detectable nitrogen cycling genes, while the remaining genomes displayed a clear functional hierarchy. Dissimilatory nitrate to nitrite reduction (DNN) was most widespread (24.2% of genomes), followed by assimilatory nitrate reduction (ANRA, 15.2%), denitrification (DNF, 12.1%), nitrogen fixation (NF, 8.7%), dissimilatory nitrite reduction to ammonium (DNRA, 8.7%), and nitrification (NIT, 0.3%) (Figure 1A). Among functionally active genomes, single-pathway specialists dominated (Figure 1B; Supplementary Note 1).

**Figure 1.**
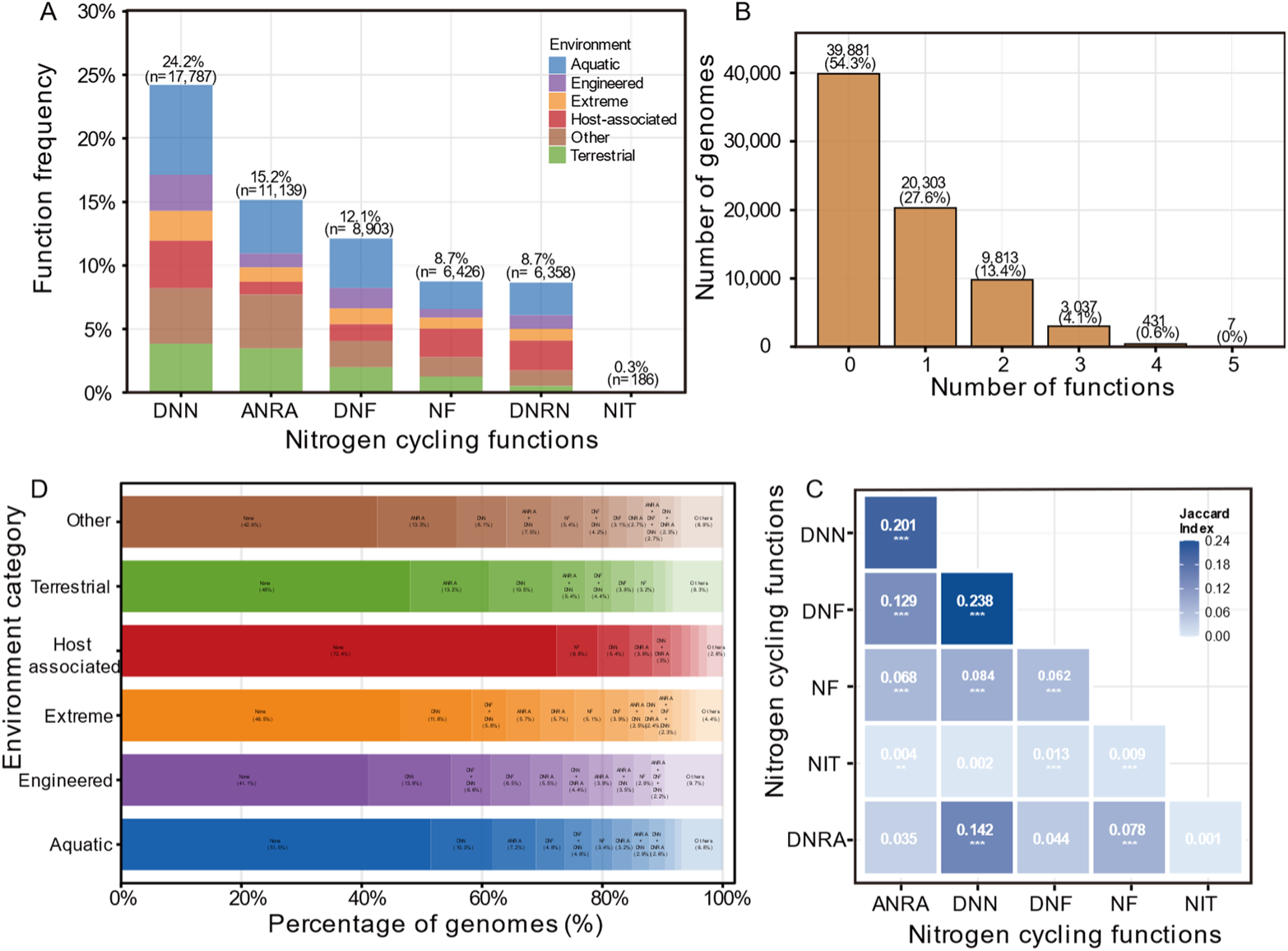
Nitrogen cycling functional distributions and co-occurrence patterns across bacterial genomes. (A) Function frequency distribution among 73,472 GTDB genomes (B) Functional richness distribution per genome. (C) Jaccard similarity matrix showing pairwise co-occurrence patterns. (D) Function combinations within genome source categories. Statistical significance: ***p<0.001, **p<0.01, *p<0.05. ANRA, assimilatory nitrate reduction; DNN, dissimilatory nitrate reduction to nitrite; DNF, denitrification; NF, nitrogen fixation; DNRA, dissimilatory nitrite reduction to ammonium; NIT, nitrification.

Functional co-occurrence analysis revealed systematic patterns (Figure 1C; Supplementary Note 1). The strongest association occurred between sequential denitrification steps (DNF-DNN, Jaccard = 0.238). The second-strongest linkage unexpectedly connected DNN with the assimilatory pathway ANRA (Jaccard = 0.201), exceeding the biochemically contiguous DNN-DNRA connection (Jaccard = 0.142).

Genome source associations revealed systematic relationships between functional capabilities and environmental context (Figure 1D; Supplementary Note 1). Aquatic isolates showed enrichment for denitrification pathways, while terrestrial isolates exhibited pronounced enrichment for assimilatory processes. Host-associated sources displayed a distinctive functional signature: despite having the highest proportion of genomes lacking nitrogen cycling functions (72.4%), they exhibited the strongest enrichment for nitrogen fixation among functionally active genomes. These distribution patterns prompted examination of taxonomic constraints on functional potential.

### Taxonomic constraints on nitrogen cycling potential vary inversely with functional prevalence

To what extent does taxonomic identity constrain nitrogen cycling functional distribution? Hierarchical variance decomposition across all six nitrogen transformation pathways revealed that taxonomic identity substantially constrains functional potential, explaining 20.0-46.1% of functional variation (Figure 2). Constraint strength exhibits a systematic hierarchy inversely correlating with functional prevalence: DNRA (46.1%, 241 genera/24 classes) > NF (44.2%, 328 genera/20 classes) > DNF (41.4%, 447 genera/16 classes) > DNN (38.6%, 653 genera/26 classes) > ANRA (36.4%, 531 genera/16 classes) > NIT (20.0%, 6 genera/3 classes; interpretation limited by extreme rarity, see Discussion).

**Figure 2.**
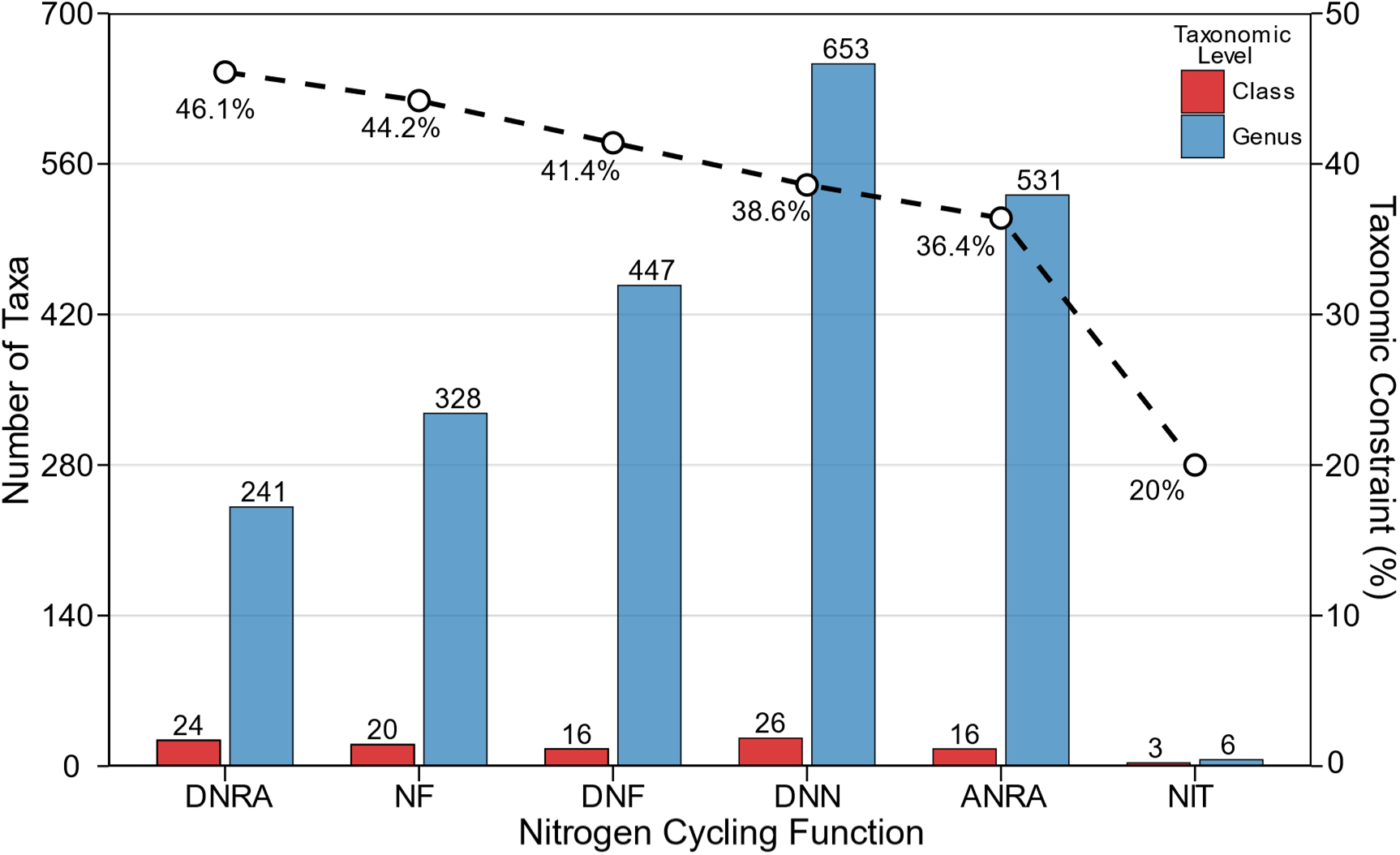
Taxonomic constraint strength and distribution patterns of nitrogen cycling functions. Bars show the number of classes (red) and genera (blue) harboring each function (left y-axis). Dashed line shows constraint strength (right y-axis, % variation explained by taxonomy).

Variance component analysis demonstrated that taxonomic constraints operate primarily at the genus level rather than class level across all functions (Supplementary Note 2). For highly constrained pathways (DNRA, NF), genus-level variance components were 3.2-3.8-fold higher than class-level components, while for weakly constrained pathways (ANRA, DNN), this ratio was 2.1-2.4-fold. These genus-level constraints raised the question: do recognizable organizational patterns emerge at broader taxonomic scales?

### Four functional archetypes organize class-level nitrogen cycling potential

Unsupervised K-means clustering based on participation rate and functional concentration across 77 bacterial classes identified four distinct functional archetypes (Functionally Inactive, Diversified, Integrated, and Specialized; ≥100 genomes each), with k = 4 selected based on maximum silhouette score (0.637) (Figure 3A).

**Figure 3.**
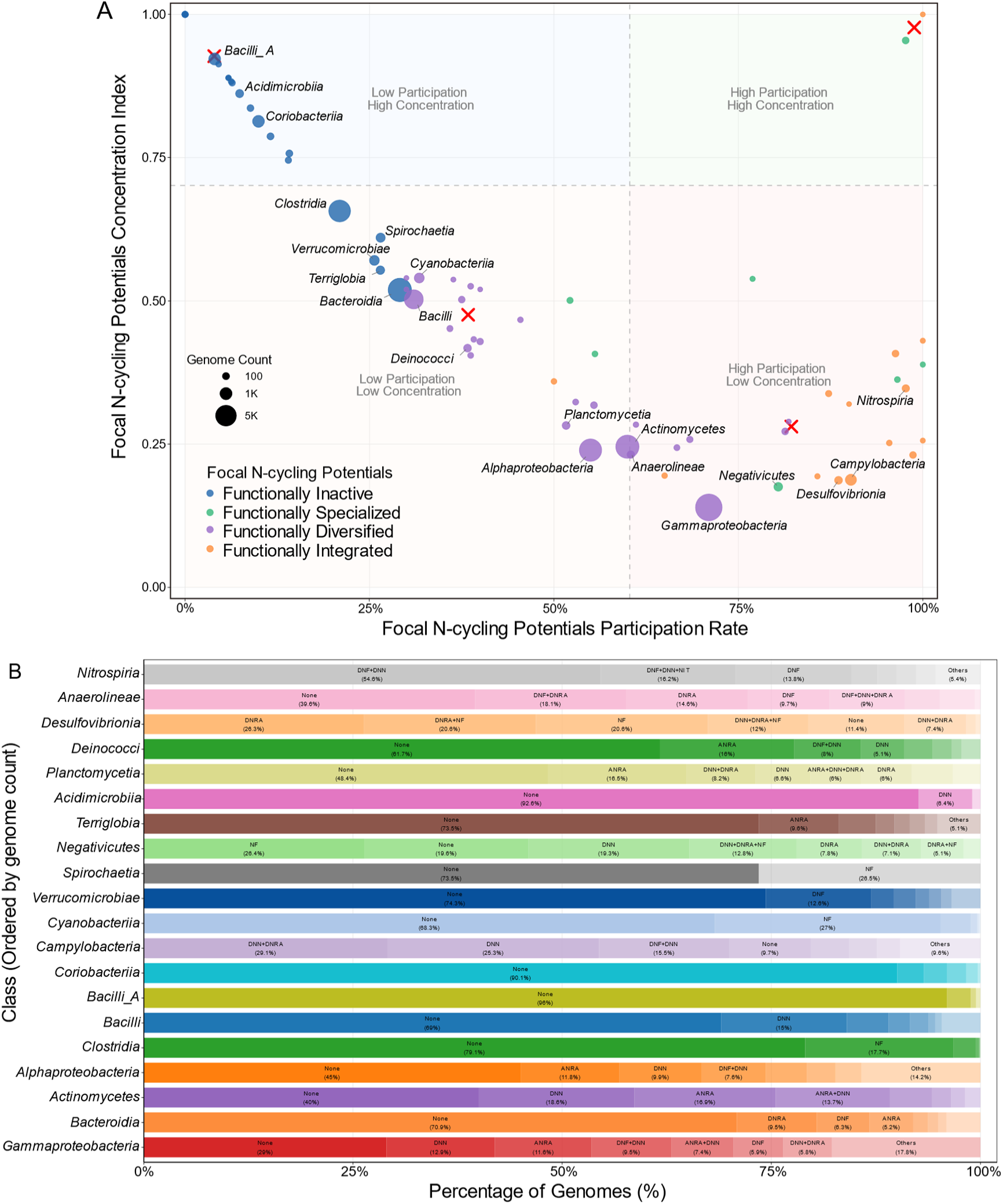
Class-level functional strategy differentiation and environmental modulation patterns. (A) Functional strategy classification of 77 bacterial classes positioned by participation rate and functional concentration index. K-means clustering (k = 4) identified four strategy archetypes. Point size represents genome count per class. (B) Environmental modulation of functional strategies. Heatmap showing functional frequency variations across major bacterial classes within six genome source categories.

Functionally Inactive dominated with 30 classes (39.0%), characterized by widespread functional absence and specialized retention when present. Bacteroidia (6,816 genomes) displayed 70.9% functional absence while active participants demonstrated focused profiles (DNRA 9.5%, DNF 6.3%, ANRA 5.2%; Supplementary Note 3). Functionally Diversified comprised 26 classes (33.8%), achieving balanced functional distribution with high participation rates. Gammaproteobacteria (9,466 genomes) exhibited 71.0% participation with balanced pathway representation (DNN 12.9%, ANRA 11.6%) and diverse functional combinations (Supplementary Note 3). Functionally Integrated encompassed 14 classes (18.2%), demonstrating coordinated functional integration. Campylobacteria (759 genomes, 90.3% participation) showed multi-functional combinations (DNN 25.3%, DNN+DNRA 29.1%) exceeding single pathways (Supplementary Note 3). Functionally Specialized included 7 classes (9.1%), exhibiting concentrated pathway investment. Cyanobacteriia (523 genomes) displayed concentrated nitrogen fixation (27.0%) despite 68.3% overall functional absence.

Genome source associations revealed stability of these taxonomy-based organizations across environments (Figure 3B and Figure S3). Classes consistently maintained archetypal characteristics across environments. Gammaproteobacteria preserved its diversified-strategy identity across all genome sources while showing systematic variation in specific function frequencies, with participation rates ranging from 57.8% to 74.2% (Supplementary Note 3).

### Five ecological strategies balancing nitrogen retention and loss exhibit strong phylogenetic conservatism

Class-level archetypes establish broad functional frameworks, but how do these constraints manifest at finer taxonomic resolution? We classified 1,281 qualified genera (≥10 genomes each) into five ecological strategy types based on nitrogen retention versus loss functional profiles (Figure 4A). N-Retention dominated with 468 genera (36.5%), characterized by pathways conserving nitrogen through assimilatory nitrate reduction (ANRA), nitrogen fixation (NF), and dissimilatory nitrate reduction to ammonium (DNRA). Multifunctional comprised 387 genera (30.2%), featuring capabilities spanning both retention and loss pathways. Non-N-Cycling included 285 genera (22.2%) lacking detectable nitrogen cycling functional markers. N-Loss encompassed 77 genera (6.0%) specializing in nitrogen removal through denitrification (DNF) and nitrification (NIT). Metabolically Versatile represented 64 genera (5.0%) exhibiting dissimilatory nitrate to nitrite reduction (DNN) without committed retention or loss pathways.

**Figure 4.**
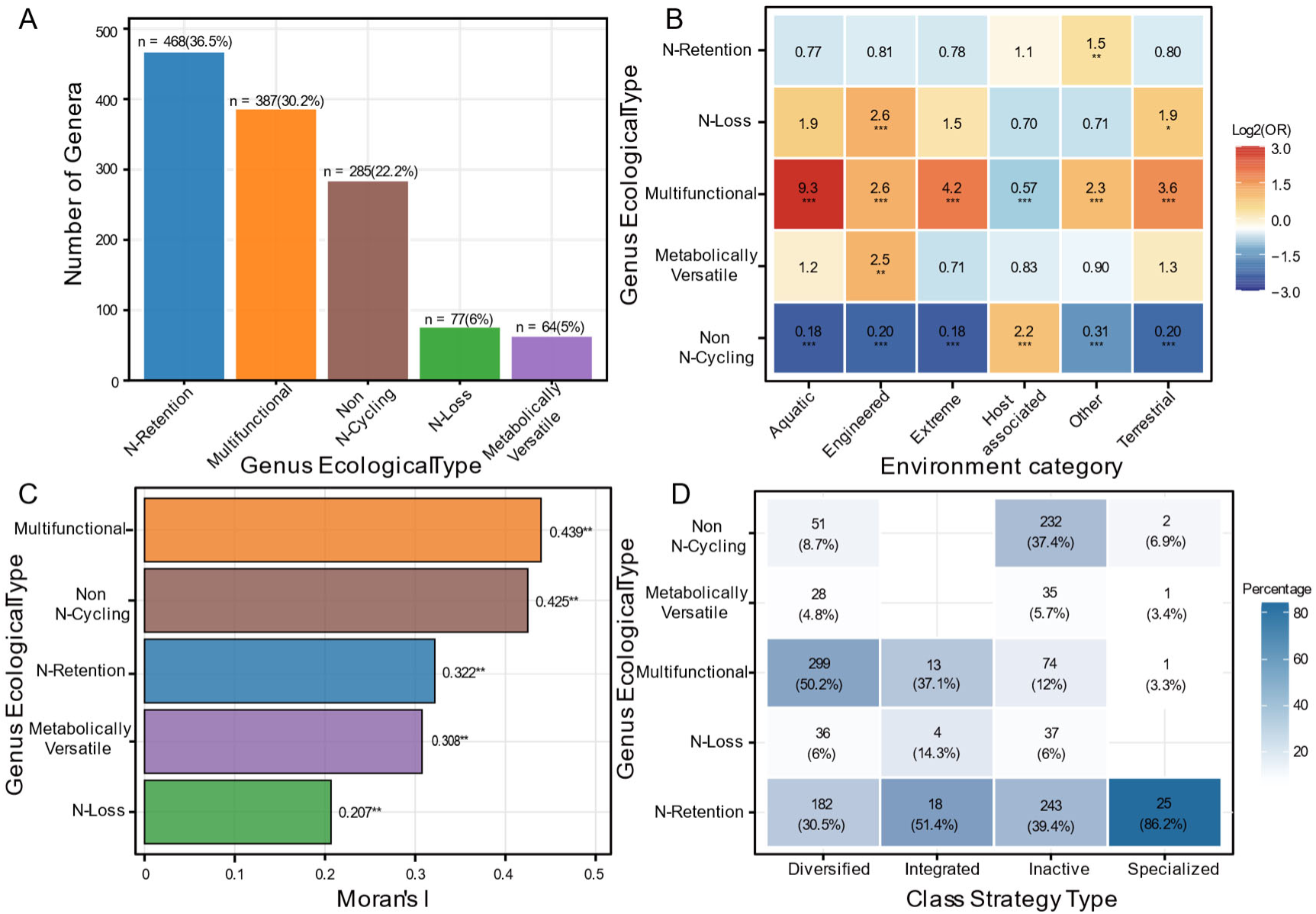
Genus-level ecological strategy classification, phylogenetic constraints, and multi-scale validation. (A) Ecological strategy type distribution across 1,281 qualified genera showing five strategy types based on nitrogen retention versus loss functional profiles. (B) Genome source-ecological strategy associations via Fisher’s exact test. Red: enrichment (OR >1.5), blue: depletion (OR <0.67). (C) Phylogenetic signal strength in ecological strategy types. Moran’s I values for five ecological strategy types. (D) Strategy-ecology association matrix demonstrating phylogenetic constraints. Percentage distribution of genus ecological types within each class strategy type. Statistical significance: ***p < 0.001, **p < 0.01, *p < 0.05.

Ecological strategies showed systematic genome source associations (Figure 4B; Supplementary Note 4). Aquatic-sourced genomes showed strong statistical association with Multifunctional genera (OR = 9.27, q < 0.001, Benjamini-Hochberg FDR correction) while showing relative depletion of Non-N-Cycling genera (OR = 0.18, q < 0.001). Host-associated genome sources showed statistical enrichment for Non-N-Cycling strategies (OR = 2.20, q < 0.001) while showing reduced representation of Multifunctional genera (OR = 0.57, q < 0.001). Of 30 genome source-strategy combinations tested, 16 achieved statistical significance (53.3%, q < 0.05).

Ecological strategies exhibited strong phylogenetic conservatism (Figure 4C and Figure S4). Moran’s I analysis revealed systematic phylogenetic structuring across all five strategy types (I = 0.207–0.439, all p < 0.001), with Multifunctional (I = 0.439) and Non-N-Cycling (I = 0.425) showing the strongest signals. This contrasted dramatically with genome source preferences (I = 0.032) and individual function frequencies (I = 0.033). The phylogenetic tree showed non-random clustering of strategies within bacterial lineages, with Non-N-Cycling genera concentrated in Clostridia and Bacilli, while Multifunctional genera enriched in Gammaproteobacteria and Alphaproteobacteria (Figure S4; Supplementary Note 4).

The hierarchical constraint framework predicts that class-level archetypes should constrain genus-level strategies. Analysis demonstrated that 78.7% of genera conformed to their class-level expectations (Figure 4D and Figure S6). Functionally Specialized classes predominantly corresponded to N-Retention genera (84.6%), while Functionally Diversified classes frequently produced Multifunctional genera (50.1%). Functionally Inactive classes exhibited bifurcated patterns between N-Retention (39.4%) and Non-N-Cycling (36.9%) types (Supplementary Note 4). The remaining 21.3% of genera deviated from class-level expectations. Complete functional profiles and ecological strategy classifications for all 1,281 qualified genera are documented in Supplementary Table S2.

### Sequence conservation and functional constraints operate as partially independent dimensions in nitrogen cycling genes

Having established hierarchical functional organization, we examined whether these macro-evolutionary patterns are reflected at the molecular level. Nucleotide diversity (π) analysis across 13 core nitrogen cycling genes addressed whether sequence conservation follows the same multi-scale hierarchy (genus > class > environment) as functional distribution, and whether gene-specific evolutionary patterns reveal mechanisms underlying taxonomic constraints.

Sequence conservation followed the predicted hierarchical gradient (Figure 5A). Across all genes, π was lowest at the genus level (mean = 0.229), intermediate at the class level (mean = 0.344), and highest when grouped by genome source environment (mean = 0.433), with all pairwise differences statistically significant (paired Wilcoxon signed-rank tests, p < 0.001). Gene-specific scale dependency revealed systematic variation, with *cnorB, nosZ*, and *napA* displaying pronounced phylogenetic structuring (environment/genus π ratios 2.22-2.28-fold).

**Figure 5.**
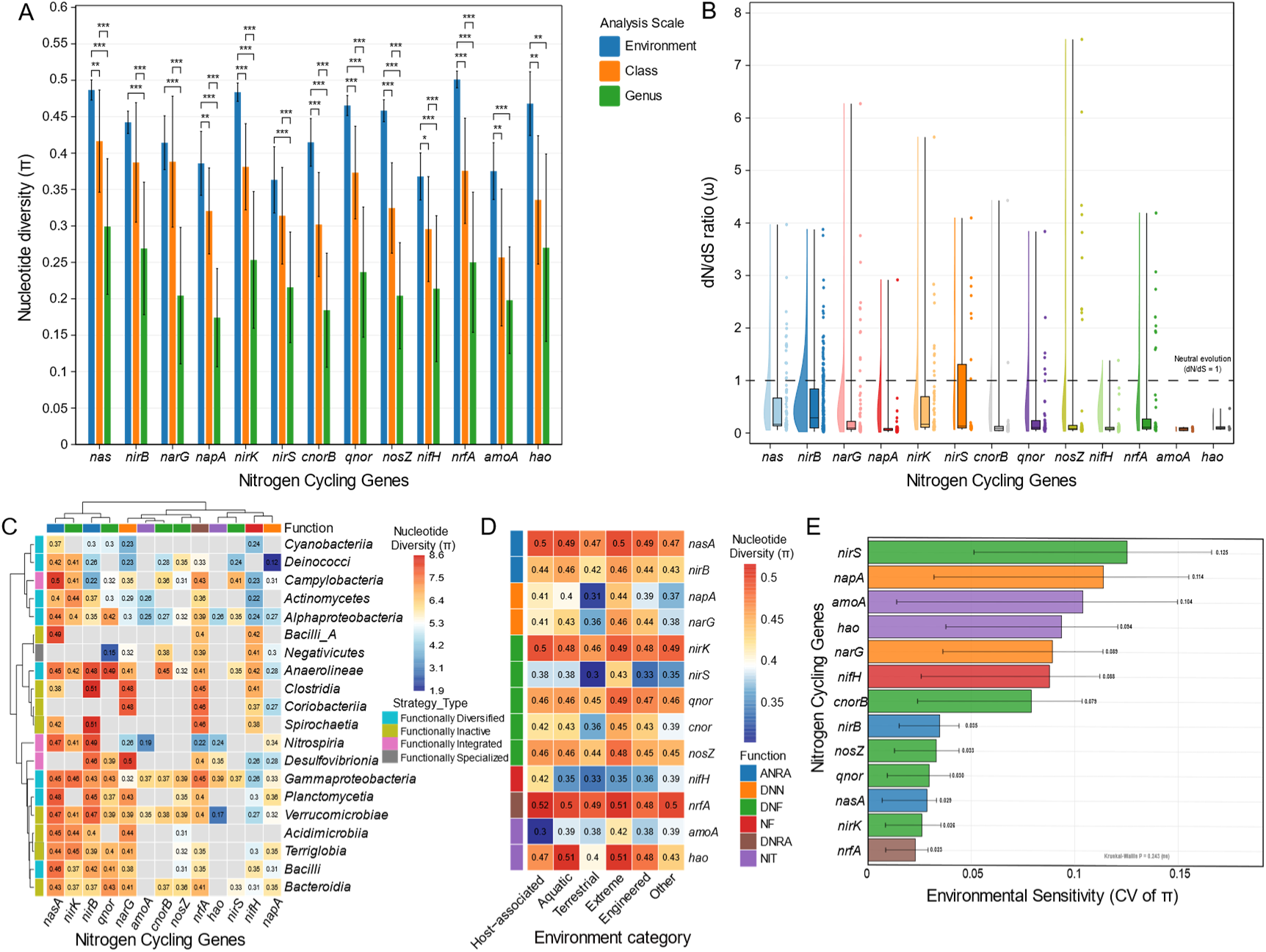
Multi-scale molecular evolutionary patterns and environmental modulation of nitrogen cycling genes. (A) Nucleotide diversity (π) for 13 nitrogen cycling genes across genome source (orange), class (blue), and genus (green) levels. Error bar show 95% confidence intervals. Kruskal-Wallis test followed by Dunn’s post-hoc comparisons, ***p<0.001; (B) Distribution of dN/dS ratios (ω) across genera with ≥20 sequences. Red dashed line indicates neutral evolution threshold (ω = 1). Genes ordered by π from (A). Detailed statistics in Supplementary Table S4. (C) Gene-class conservation matrix showing nucleotide diversity (π) across 20 major bacterial classes and 13 nitrogen cycling genes. Color scale: blue (low) to red (high conservation). Gray cells indicate gene absence. (D) Environmental sensitivity of nitrogen cycling genes quantified as nucleotide diversity (π) across six genome source categories. Higher values indicate greater sequence diversity. Gray cells: insufficient data. (E) Genome source sensitivity measured as coefficient of variation (CV) of π values across the six environmental categories. Error bars show 95% confidence intervals.

dN/dS ratio (ω) analysis revealed inconsistent relationships between functional constraint strength and sequence conservation. Four distinct patterns emerged (Figure 5B, Supplementary Table S4). First, congruent constraint-conservation relationships characterized genes from pathways at constraint extremes. Nitrogenase reductase *nifH* from the highly constrained nitrogen fixation pathway (44.2% functional constraint) exhibited correspondingly strong sequence conservation (π = 0.21 ± 0.10, median ω = 0.081), with 91.9% of genera under strong purifying selection (ω < 0.3). Conversely, assimilatory nitrate reductase *nasA* and nitrite reductase *nirB* from the weakly constrained ANRA pathway (36.4% functional constraint) displayed relaxed sequence conservation (π = 0.27-0.30) with more permissive selection regimes.

Second, pathway-level constraint-conservation decoupling manifested as under-conservation in cytochrome c nitrite reductase *nrfA*. Despite being the core enzyme of the most functionally constrained pathway (DNRA, 46.1%), *nrfA* exhibited only moderate nucleotide diversity at the genus level (π = 0.25 ± 0.096, ranking 9th among 13 genes) with a bimodal dN/dS distribution. While 74.5% of 47 analyzed genera showed strong purifying selection (ω < 0.3, median 0.108), 19.1% exhibited relaxed or positive selection (ω ≥ 1.0)—approximately ten-fold higher than *nifH* (1.6%). Multi-scale nucleotide diversity analysis revealed that this heterogeneity is structured by phylogenetic relationships rather than ecological contexts. Environmental analysis showed that *nrfA* maintained consistent π values across all six environmental categories (range: 0.485-0.516, CV = 0.023; Figure 5D-E). In contrast, class-level π values varied substantially across bacterial lineages (range: 0.390-0.463; Figure 5C; Supplementary Note 5). The nine high-ω genera maintain high DNRA functional prevalence (mean 70.1%) and span seven phylogenetically distinct classes (Supplementary Note 5).

Third, pathway-level constraint-conservation decoupling manifested as over-conservation in periplasmic nitrate reductase *napA*. Despite originating from the moderately constrained DNN pathway (38.6%), *napA* displayed the lowest nucleotide diversity (π = 0.17 ± 0.067) and median ω (0.057) among all nitrogen cycling genes, with 93.5% of genera under strong purifying selection, significantly exceeding the more prevalent membrane-bound nitrate reductase *narG* (π = 0.20, median ω = 0.096, 75% strong purifying selection; p < 0.001). Multi-scale analysis revealed contrasting patterns: class-level analysis showed variable conservation across lineages (Figure 5C; Supplementary Note 5), while environmental analysis revealed substantial π variation (range: 0.312 in terrestrial to 0.440 in extreme environments, CV = 0.11; Figure 5D-E).

Fourth, gene-specific heterogeneity within pathways rivaled variation between pathways with divergent functional constraints. In denitrification (DNF, 41.4% constraint), gene-specific nucleotide diversity at the genus level ranged from 0.18 (*cnorB*) to 0.25 (*nirK*), a 1.39-fold range approaching the difference between *nifH* (0.21) and *nirB* (0.27, 1.29-fold) from pathways with substantially divergent functional constraints (44.2% vs 36.4%). Nitric oxide reductase *cnorB* exhibited the highest conservation within denitrification (median ω = 0.089, 97.2% genera ω < 0.3). The evolutionarily unrelated nitrite reductases *nirS* and *nirK* showed significant differences in genus-level conservation (π: 0.22 vs 0.25, p < 0.001; high-ω genera: 30.8% vs 17.7%). Environmental sensitivity analysis reinforced this gene-specific autonomy: *nirS* displayed dramatic environmental modulation (range: 0.302 in terrestrial to 0.432 in extreme environments, CV = 0.13), while *nirK* exhibited remarkable environmental stability (CV = 0.026), revealing a striking 4.8-fold difference in environmental sensitivity between genes catalyzing identical reactions (Figure 5D-E; Supplementary Note 5).

## Discussion

Our multi-scale analysis reveals that taxonomic constraints on nitrogen cycling operate through hierarchical mechanisms spanning functional distribution, ecological strategy organization, and molecular evolution. These constraints vary systematically across pathways and biological scales. We examine how pathway complexity, phylogenetic inheritance, and gene-specific structural factors interact to produce these patterns, revealing that functional and molecular constraints operate as partially independent evolutionary dimensions.

The inverse relationship between functional constraint strength and prevalence reflects fundamental evolutionary dynamics governing nitrogen cycling gene distribution. Highly constrained pathways like DNRA and nitrogen fixation involve complex multi-component enzyme systems requiring coordinated expression, cofactor biosynthesis, and integration with cellular metabolism, creating barriers to horizontal transfer. In contrast, weakly constrained pathways such as ANRA and DNN possess simpler, modular genetic architectures that facilitate dissemination across diverse lineages via HGT (10). This mechanistic basis explains why certain nitrogen cycling capabilities cluster within specific taxonomic lineages while others show broader phylogenetic distributions (6). The dominance of genus-level over class-level variance components indicates that functional coherence emerges most strongly at fine taxonomic scales reflecting recent evolutionary divergence and shared metabolic inheritance patterns (38).

These quantified constraint strengths establish that taxonomic identity provides informative but probabilistic associations with functional potential rather than deterministic predictions, forming the basis for systematic functional inference across taxonomic scales. While genus-level taxonomic identity explains up to 46% of functional variation for highly constrained pathways, the remaining unexplained variation reflects multiple evolutionary processes including HGT events, lineage-specific gene loss, and environmental adaptation within taxonomic lineages (12). This finding underscores that taxonomic constraint represents a probabilistic, not deterministic, predictive framework that must account for evolutionary flexibility within taxonomic boundaries. The interpretation for NIT (20.0% constraint) requires particular caution due to severe underrepresentation of nitrifying organisms in current genome databases (39), highlighting database-dependent limitations. These parameters establish quantitative foundations for taxonomically-informed inference and provide the framework for examining multi-scale functional organization.

The emergence of four discrete archetypes—rather than continuous variation—indicates that taxonomic constraints channel functional organization into specific regions of evolutionary strategy space. The four archetypes represent distinct solutions to metabolic resource allocation. Functionally Inactive classes exemplify genome reduction where metabolic investments are selectively minimized when alternative nitrogen sources are available (40). Functionally Diversified classes maintain diverse functional potential for success in variable environments. Functionally Integrated classes reflect evolutionary optimization for specific biogeochemical niches requiring coordinated nitrogen transformations. Functionally Specialized classes demonstrate deep metabolic investment in specific transformations where strong taxonomic constraints confer competitive advantages in specialized niches. These archetypal patterns likely reflect fundamental trade-offs in metabolic resource allocation under different evolutionary pressures, where taxonomic constraints shape the accessible solution space for nitrogen cycling organization.

Genome source associations revealed the stability of these taxonomy-based functional organizations while revealing how ecological context modulates their expression. Classes consistently maintain their archetypal characteristics across environments, indicating that taxonomic identity provides the fundamental framework for functional organization. This environmental modulation occurs within the boundaries set by taxonomic constraints, demonstrating that while ecological pressures shape specific functional manifestations, the underlying organizational principles remain taxonomically determined. These patterns establish that class-level functional archetypes represent stable taxonomic characteristics that provide a robust framework for predicting nitrogen cycling capabilities across diverse environmental contexts.

The dominance of N-Retention (36.5%) and Multifunctional (30.2%) strategies, comprising 66.7% of genera, reflects fundamental ecological trade-offs in bacterial nitrogen metabolism. N-Retention represents a conservative strategy consistent with specialist frameworks, while Multifunctional genera maintain capabilities across both retention and loss pathways, providing metabolic versatility for adaptation to variable nitrogen availability. The strong aquatic-Multifunctional association (OR = 9.27) is consistent with variable redox conditions in aquatic environments where metabolic flexibility provides advantages (11), while Non-N-Cycling enrichment in host-associated sources (OR = 2.20) reflects genome streamlining principles in stable, nutrient-rich environments (41).

Ecological strategies showed much stronger phylogenetic signals than individual functional components, suggesting that strategy-level classifications capture phylogenetic constraints more effectively than individual gene distributions. This disparity reflects evolutionary dynamics where individual genes may be gained or lost through HGT, while complete multi-gene functional modules that constitute ecological strategies are inherited and maintained as coherent evolutionary units (42), consistent with greater phylogenetic conservation of complex traits (7, 38). The 78.7% cross-scale consistency between class-level archetypes and genus-level strategies supports hierarchical constraint organization, where broader taxonomic identity constrains the accessible strategy space while permitting lineage-specific refinements. The remaining 21.3% deviations reflect evolutionary flexibility where ecological pressures and HGT enable departures from class-level predictions while maintaining overall taxonomic coherence (12).

Four distinct constraint-conservation patterns—congruence (*nifH*), under-conservation (*nrfA*), over-conservation (*napA*), and within-pathway heterogeneity (denitrification genes)—reveal that sequence evolution is governed by selective pressures operating hierarchically across biological scales. The congruent pattern reflects the expected relationship where complex, multi-component enzyme systems subject to strong vertical inheritance maintain stringent sequence fidelity, while simpler, modular pathways permit greater evolutionary flexibility. The *nrfA* under-conservation pattern, where 19.1% of genera exhibit relaxed or positive selection despite maintaining high functional prevalence (mean 70.1%), indicates that the bimodal selection distribution reflects lineage-specific variation in physiological integration within distinct metabolic architectures (43–45). Given that DNRA is deployed across diverse regulatory contexts and electron donor regimes in different bacterial lineages, these patterns suggest that lineage-specific optimization of *nrfA* sequences for integration within divergent metabolic networks necessitates particular sequence configurations while preserving fundamental nitrite reduction chemistry. The *napA* over-conservation pattern becomes interpretable through its unique physiological deployment: unlike *narG*’s function during strict anaerobic respiration, *napA* enables periplasmic nitrate scavenging under microoxic-to-oxic conditions (46, 47), where maintenance of electron transfer efficiency across fluctuating redox states likely imposes stringent structural requirements. The within-pathway heterogeneity exemplified by *nirS* versus *nirK* aligns with observations that *nirK* can function independently while *nirS* requires additional gene products (48), potentially accounting for their divergent environmental responsiveness.

Our results establish that taxonomic identity constrains nitrogen cycling through hierarchical mechanisms at functional, ecological, and molecular scales. Critically, these constraints operate as partially independent dimensions: pathway-level functional distributions—determined by the complexity of multi-gene systems and horizontal transfer susceptibility—do not predict gene-level sequence conservation patterns. This independence is revealed through systematic deviations where functionally constrained pathways show relaxed sequence conservation while moderately constrained pathways exhibit exceptional conservation, reflecting gene-specific factors including protein structural features, cofactor coordination requirements, and physiological deployment contexts. The hierarchical organization from class-level archetypes through genus-level strategies provides a multi-scale framework for understanding how taxonomic constraints and gene-specific factors jointly shape nitrogen cycling diversity. This framework enables probabilistic functional prediction from taxonomic composition, with applications in amplicon-based community analysis, targeted cultivation strategies, and biogeochemical modeling. The quantitative baselines for 1,281 genera represent an expandable foundation that will improve as genome databases grow. Integration with transcriptomic and environmental data represents the next step for translating genomic potential into activity predictions across dynamic conditions.

## Data availability

Bacterial genome sequences analyzed in this study were obtained from the Genome Taxonomy Database (GTDB) release 220 (https://gtdb.ecogenomic.org/). Environmental metadata were retrieved from NCBI BioSample database (https://www.ncbi.nlm.nih.gov/biosample/). Functional annotation was performed using publicly available databases: KEGG (https://www.genome.jp/kegg/), RAST server (https://rast.nmpdr.org/), and NCycDB (http://ncycdb.ncbr.mssm.edu/). All data generated in this study, including genus- and class-level functional reference baselines, ecological strategy classifications, and statistical analyses, are provided in the Supplementary Materials accompanying this article.

## Acknowledgements

This work was supported by the National Key Research and Development Program of China (2024YFD1701002), the National Natural Science Foundation of China (32400085), the Postdoctoral Fellowship Program of CPSF (GZC20231127), the National Natural Science Foundation of China (42507183), the Natural Science Foundation of Jiangxi Province (20252BAC200363), and the high-performance computing platform of Bioinformatics Center, Nanjing Agricultural University.

